# Improving access to endogenous DNA in ancient bone and teeth

**DOI:** 10.1101/014985

**Authors:** Peter B. Damgaard, Ashot Margaryan, Hannes Schroeder, Ludovic Orlando, Eske Willerslev, Morten E. Allentoft

## Abstract

Poor DNA preservation is the most limiting factor in ancient genomic research. In the vast majority of ancient bones and teeth, endogenous DNA molecules only represent a minor fraction of the whole DNA extract, rendering traditional shot-gun sequencing approaches cost-ineffective for whole-genome characterization. Based on ancient human bone samples from temperate and tropical environments, we show that an initial EDTA-based enzymatic ‘pre-digestion’ of powdered bone increases the proportion of endogenous DNA several fold. By performing the pre-digestion step between 30 min and 6 hours on five bones, we identify the optimal pre-digestion time and document an average increase of 2.7 times in the endogenous DNA fraction after 1 hour of pre-digestion. With longer pre-digestion times, the increase is asymptotic while molecular complexity decreases. We repeated the experiment with n=21 and t=15-30’, and document a significant increase in endogenous DNA content (one-sided paired t-test: p=0.009). We advocate the implementation of a short pre-digestion step as a standard procedure in ancient DNA extractions from bone material. Finally, we demonstrate on 14 ancient teeth that crushed cementum of the roots contains up to 14 times more endogenous DNA than the dentine. Our presented methodological guidelines considerably advance the ability to characterize ancient genomes.

## Introduction

With the introduction of next-generation sequencing (NGS) technology, ancient DNA (aDNA) research has over the last decade advanced from retrieving short segments of mostly mitochondrial DNA (mtDNA) to the assembly of whole nuclear genomes (reviewed in (Der Sarkissian et al., 2015)). Amongst many highlights such paleogenomic research has for example provided documentation for Late Pleistocene admixture between Neanderthals and anatomically modern humans (Green et al., 2010; Prüfer et al., 2014), has assisted in the discovery and description of a previously unknown hominin, the Denisovan (Krause et al., 2010; Meyer et al., 2012; Reich et al., 2010), yielded detailed insights into early human colonisation of the Americas and the Arctic (Raghavan et al., 2013, 2014b; Rasmussen et al., 2010, 2014) and clarified equine evolution by sequencing the complete genome of a 700,000 years old horse (Orlando et al., 2013).

Although with some exceptions (Fu et al., 2014; Seguin-Orlando et al., 2014) it has been common to all of the ancient hominin samples from which ancient whole-genomes have been sequenced (Gamba et al., 2014; Keller et al., 2012; Lazaridis et al., 2014; Meyer et al., 2012; Olalde et al., 2014; Prüfer et al., 2014; Raghavan et al., 2013; Rasmussen et al., 2010, 2014; Skoglund et al., 2014b) that they displayed an exceptional biomolecule preservation, in particular as most of the DNA sequencing libraries contained a high proportion (28.2 % – 70 %) of authentic human DNA molecules, hereafter referred to as the ′endogenous′ DNA content. However, in the vast majority of DNA extracts from ancient bones and teeth the endogenous DNA content is less than 1% (Carpenter et al., 2013; Rizzi et al., 2012) with environmental microbial DNA being highly abundant (Poinar et al., 2006; Sarkissian et al., 2014). Whole genomes, or genome-wide information, are preferentially sequenced using Next Generation shotgun Sequencing, but when the target molecules represent such a minute fraction this is either not feasible or, at best, extremely expensive. Although novel genomic capture methods can be used to enrich for the target DNA (Carpenter et al., 2013), most ancient samples will remain unsuitable for whole-genome characterization. Therefore a common approach in paleogenomic research is to extract and shotgun sequence a large number of samples in an initial screening phase in order to identify a few good candidate samples. This approach is time-consuming and expensive, and requires the destruction of many samples that will not be amenable to genome analyses. Thus, the field of paleogenomics would benefit greatly from methodological advances that could increase the endogenous DNA fraction during DNA extraction. In this study we present two such advances.

Although the biochemical processes driving DNA preservation in bone are not yet fully understood, it has been shown that ancient DNA (aDNA) is preserved both in association with the bone minerals, hydroxy-apatite aggregates, and within the organic composite, i.e. the collagen fibrils (Campos et al., 2012; Schwarz et al., 2009). During decomposition the bone structure degrades which increases the porosity and total surface area of the bone (Campos et al., 2012; Jans, 2008). Although microorganism colonization will likely be heterogenous within a bone (Collins et al., 2002), we would expect that during the digestion of bone material in the first step of an aDNA extraction, surface contaminants will be released into solution first, regardless of their exact location. This is in contrast to the endogenous DNA which might be better protected within the bone′s microniches as previously suggested (Ginolhac et al., 2012). We therefore hypothesized that treating the grinded bone material with a digestion buffer for a short period of time (a “pre-digestion”) would remove a fraction of the exogenous non-target DNA and thereby enrich the DNA extract for endogenous DNA. Similarly, we hypothesized that modern human DNA contamination, deposited on the bone surface during recent handling, would also be preferentially removed with such pre-digestions. Indeed, higher endogenous DNA fractions were recently observed on second extractions undertaken on remaining bone pellets that had not been fully dissolved after 24 hours of incubation in a digestion buffer (Ginolhac et al., 2012; Orlando et al., 2011; Sarkissian et al., 2014). These promising observations provided an impetus for a more systematic assessment of this phenomenon in order to validate the potential for implementing an initial chemical cleaning into standardized aDNA extraction protocols.

We used Next Generation shotgun sequencing to monitor the changes in endogenous DNA content and sequence complexity in DNA extracts from the same bone treated with varying pre-digestion times. Although specific sample substrate and its preservation state impacts on the dissolution rate during the treatment, we aspired to produce a general recommendation regarding an optimal pre-digestion time. Four bones from Easter Island (post-1200 AD), and one bone from Copenhagen, Denmark (18^th^ century) were included in this initial experiment. Additionally a total of 21 bones from Easter Island (post-1200 AD), Hungary (Bronze-Age 2000-1500 BC), and Guadeloupe (400-1400 AD) were used in a follow-up experiment to confirm the efficiency and test the significance of the general improvement when applying brief pre-digestions (i.e. 15-30 minutes).

Teeth roots have been demonstrated as an excellent resource for aDNA (Adler et al., 2011; Higgins and Austin, 2013; Higgins et al., 2013) but a systematic comparison of the endogenous DNA content in the root cementum and root dentine is lacking. It has been shown that while the nuclear DNA concentrations decline drastically in the dentine throughout the life of an individual (Trivedi et al., 2002), levels of nucleated cells in the apical cementum layer are unaffected by age (Higgins et al., 2013). Moreover, a quantitative PCR approach determined that in ancient teeth the concentration of human mtDNA is generally elevated in the cementum compared to the dentine (Adler et al., 2011). However, the proportion of endogenous DNA may still be low as the cementum layer is exposed at the root surface and thus potentially more affected by microbial colonization than dentine. To investigate this we extracted DNA and used shotgun sequencing to estimate the endogenous DNA proportions in crushed root surface (containing mainly the cementum layer), and the deeper parts of the root (containing mainly dentine) from 14 ancient teeth (Table 1), from Denmark (18^th^ century and Iron Age c. 100 AD), Easter Island (post-1200 AD), and Greenland (c. 1100 AD).

**Table 1:**
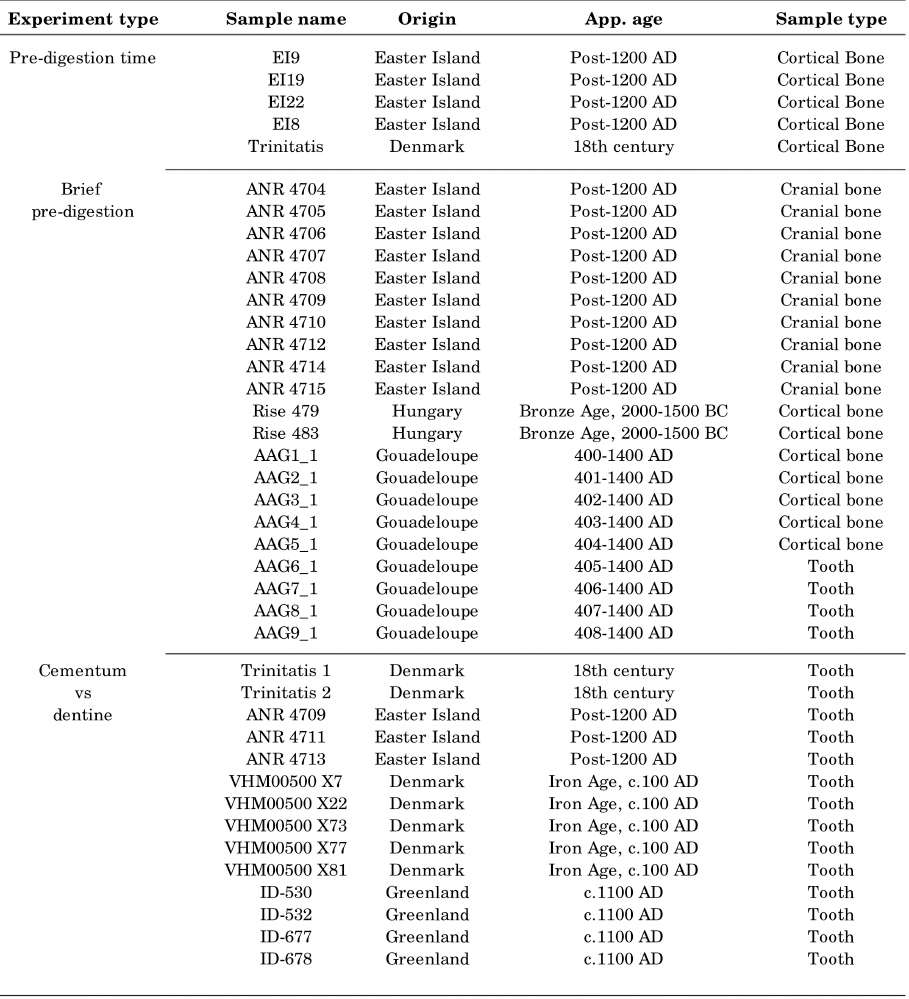
Sample information.

## Materials and methods

All the laboratory work was performed in the dedicated clean laboratory facilities at Centre for GeoGenetics, Natural History Museum, University of Copenhagen, according to strict aDNA standards (Cooper and Poinar, 2000; Gilbert et al., 2005b; Willerslev and Cooper, 2005).

### Samples

A total of 26 ancient human bones from various archaeological contexts spanning tropical and temperate enviroments were included in the pre-digestion experiments. An additional 14 ancient teeth were used in the comparison between DNA extracted from dentine and cementum respectively. All relevant information about the samples are provided in Table 1.

### DNA extraction

All bones were cleaned on the surface using a cloth with 10% hypochlorite, followed by removal of the outer surface using a scalpel or a sterile drill bit. Five bones were selected for the initial experiment aiming to optimize pre-digestion time (Table 1). Cortical bone mass was drilled and homogenized, and 400 mg of bone powder was transferred to each of six 15 mL Falcon tubes (labelled A-F), yielding 30 extractions in total. To counteract the effect of granular convection by which the smallest bone particles end up in the first tubes, we attempted to homogenize the powder between each transfer. The bone powder was subjected to a digestion buffer containing 4.7 mL 0.5 M EDTA, 50 µL recombinant Proteinase K, and 250 µL 10% N-Laurylsarcosyl and incubated at 50ºC, slightly higher than recommended in (Orlando et al., 2013). At 30 minutes, extraction A was centrifuged and the supernatant removed. An identical digestion buffer was then transferred to the undigested and sedimented bone powder, and the sample was vortexed and returned to incubation. Similarly, pre-digest supernatants were removed for extractions B-E at time points: 1 hour, 2 hours, 3 hours, 6 hours respectively. No pre-digest supernatant was removed for extraction F. At 24 hours all digestions were stopped. The samples were then centrifuged and the supernatants transferred to new tubes for DNA extraction. The experimetal setup is illustrated in Figure 1.

**Figure 1:**
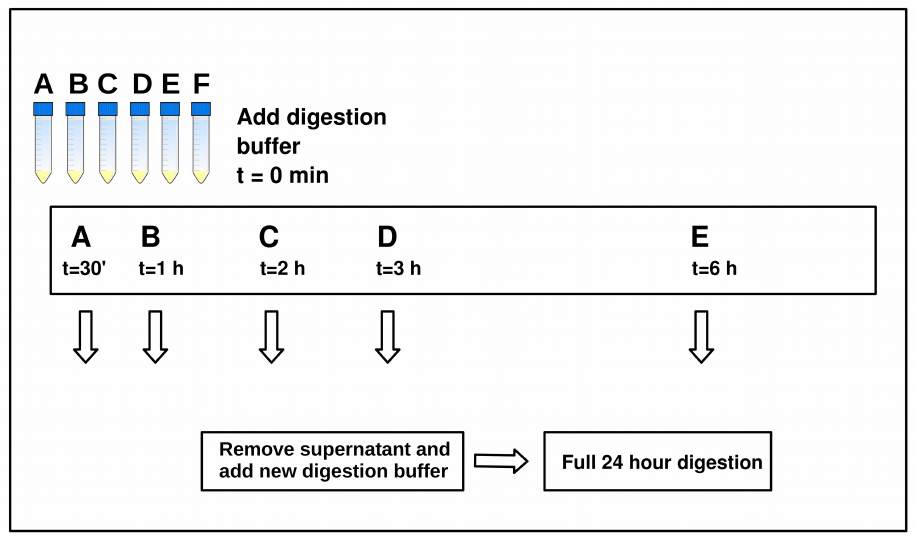
Illustration of the pre-digestion length experiment performed on 5 different bones. For each bone, approximately 2.5 g of bone powder was homogenized and 400 mg distributed into 6 tubes labelled A-F. A digestion buffer was added to all samples at t=0, samples were vortexed, and left on rotator at 50 °C. At t=30 minutes, the pre-digestion was removed from sample A and a new digestion buffer was added; similarly, pre-digestions were removed from samples B-E at respectively 1 hour, 2 hours, 3 hours, 6 hours. No pre-digestion was removed from sample F.

Guanidinum thiocyanate-based binding buffers (Orlando et al., 2013) were used when extracting the DNA from the digests. The buffer was prepared by mixing 118.2 g Guanidinium Thiocyanate with 10 mL Tris 1M, 1 mL NaCl 5M, 8mL EDTA 0.5M, 1 g N-Lauryl-Sarcosyl and water to a total volume of 200 mL. 20 mL of the binding buffer was transferred to each sample and left rotating for 3 hours with silica powder in solution to bind the DNA. After DNA-binding, the silica was centrifuged and washed twice with 80% cold ethanol, and the DNA eluted in 80 µl EB Buffer (Qiagen). The DNA concentration in all final extracts was measured using Qubit® Fluorometric Quantitation (Life Technologies, Grand Island, NY).

Following this initial experiment, we tested the improvement consistency with short pre-digestion times on ancient and historical bone samples from Easter Island, Guadeloupe and Hungary (Table 1). Each drilled sample was homogenized and split into two equal amounts, one that was extracted with 15 or 30 minutes of pre-digestion treatment, and one that was not pre-digested. The extraction procedure was similar to the one described above.

We used 14 ancient teeth (Table 1) to compare the endogenous DNA content of root surface (primarily cementum) against the inner root core (primarily dentine). First, the outermost surface of the teeth was removed mechanically with a drill-bit, as is standard procedure to exclude the most obvious surface contamination. Then each tooth was split with a cutting disk into two pieces (the crown and the root) on the transverse plane. The dentine was then drilled out of the root and transferred to a sterile tube, leaving a hollow root “cap” as illustrated in Figure 5. This remaining root cap, consisting mainly of the cementum layer, was then crushed with a mortar or drilled into smaller pieces before being transferred to a sterile tube. The two fractions from each tooth root (dentine-enriched and cementum-enriched, respectively) were then extracted separately as above, but without pre-digestion.

### Library preparation and sequencing

Blunt-end based Illumina sequencing libraries were built following guidelines previously outlined (Orlando et al., 2013), using the NEBNext® DNA Library Prep Master Mix Set E6070 (New England Biolabs Inc., Manual Version 2.1 (Briggs et al., 2007)). The libraries were amplified using a two-round PCR setup (Meyer and Kircher, 2010a; Rasmussen et al., 2010) with primers containing a 6 bp known index sequence. Details on the library preparation and PCR amplification conditions can be found in Supplementary Information Methods S1.

The amplified libraries were quantified on an Agilent 2200 TapeStation (Agilent Technologies, Palo Alto, CA, USA) or an Agilent Bioanalyzer 2100. The library pools were sequenced (100 bp, single read) at the Danish National High-throughput DNA Sequencing Centre. Basecalling and sequence sorting by sample-specific indexes was performed by the Sequencing Centre using CASAVA v.1.8.2.

### Data analyses

All reads were trimmed for adapter sequences using AdapterRemoval 1.5.2 (Lindgreen, 2012), with a minimal read length of 30 bp. The trimmed sequences were mapped against the human reference genome Hg19, HS Build37.1, using bwa (Li and Durbin, 2009) with the *samse* function using standard parameters except that seeding was disabled, following the recommendations from (Schubert et al., 2012). We used all sequences mapping uniquely to the human reference genome, then removed duplicates from the output bam file using the *rmdup* function in samtools (Li et al., 2009). The relevant summary statistics (Supplementary Tables S1-S3) used to estimate the endogenous DNA fraction (fraction of uniquely mapped human sequences divided by the total number of sequences passing trimming) and sequence clonality (proportion of duplicate human sequences), were extracted with a custom Perl script. The clonality of each library will increase with increased sequencing depth, implying that the overall sequencing efficiency (fraction of non-duplicated endogenous DNA sequences divided by total sequences) decreases. Hence we calculated the sequencing efficiency by randomly down-sampling the raw fastq files in order to match the smallest number of sequences per bone. This allowed for a direct comparison of sequencing efficiency within each experiment.

We also investigated the data for signatures of DNA damage. This was done in part to confirm that the profiled human DNA was not modern contamination, and in part to measure if the pre-digestion treatment would result in any obvious compositional biases in the DNA, or damage it further.

Based on the sequence length distributions of the sequences identified as human, we estimated the decay constant *k* (representing the fraction of broken bonds in the DNA backbones) and average DNA fragment length in the extract (1/*k*), as previously described (Allentoft et al., 2012; Deagle et al., 2006). A large *k* value reflects a pronounced exponential accumulation of small DNA fragments as a consequence of *post mortem* DNA breakage - a signature of highly degraded DNA. Following the approach described in (Allentoft et al., 2012), we investigated only the declining part of the distribution for each sample (40-90 bp) since the ends of the distribution are biased respectively by poor recovery of < 40 bp fragments during the DNA extractions (and the library building process), and the accumulation of >100 bp reads sequenced to the maximum length on the Illumina platform.

Using the Bayesian approach implemented in mapDamage 2.0 (Jónsson et al., 2013) with standard parameters we estimated the position-specific cytosine deamination probability (δs) as well as the probability of a base being positioned within a single-stranded overhang (λ), which thus relates directly to the average length of the overhangs, reflecting *post mortem* damage of aDNA (Briggs et al., 2007). In order to increase the accuracy of the damage estimates performed on the ancient human DNA fractions, these were based on the total mapped datasets and not the downsampled files. Outputs from mapDamage 2.0 were analysed and plotted with R (Team and others, 2012).

Sequence quality control statistics were generated using fastqc (Andrews and others, 2010) on the retrieved human sequences from each (not downsampled) file, as well as the total sequences of the library (downsampled human + non-human sequences), and were used to check for abnormalities, and particularly to investigate for potential changes in GC-content following pre-digestion. The results were plotted with R (Team and others, 2012).

Despite the implementation of strict aDNA protocols, it is difficult to completely avoid contamination from modern DNA when working with ancient human material - in particularly when dealing with samples that have been handled previously during excavation and while stored at museum collections (Allentoft, 2013). Therefore it was important to establish that any potential increase in endogenous DNA content following our treatments was not caused by, or accompanied by, increased DNA contamination. Accurate estimates of contamination levels require large amounts of genomic data, so we here restricted this analysis to samples with a mitochondrial genome coverage above five times (5X). We used contamMix (Fu et al., 2013) to estimate the level of human DNA contamination in the mtDNA sequences. This method compares for each individual the mapping affinities of its mtDNA sequences to its own consensus mitogenome sequence, relative to the mapping affinity of its mtDNA sequences to a dataset of potential contaminants represented by 311 mitogenomes from worldwide populations. The mitogenome consensus sequences were made using the samtools *mpileup* function (Li et al., 2009) and filtering the variant list outputted with bcftools with a script previously used in (Jacobsen et al., 2014) selecting only bases with a coverage > 5X and > 50% concordance between bases, in order not to incorporate bases with only limited depth-of-coverage and sequencing errors resulting from DNA damage misincorportations.

## Results

### Effects of pre-digestion time

The final digested solutions displayed a spectrum of colors, in which long pre-digestion times resulted in lighter colors of the final digest. We interpret this as an effect of removal of dirt and contaminant particles followed by intrinsic ion dissolution. This observation is accompanied by a decline in total DNA concentrations in the extracts as a function of pre-digestion time (Supplementary Fig. S1). An exponential decline model fits the DNA concentration values as a function of pre-digestion time well in three cases (R² values between 0.72 and 0.99, respective p values 2e-7 and 0 (rounded to seven decimals)), and modestly in two cases (R² values of 0.40 and 0.63 respectively), see Supplementary Fig. S1.

By aligning on average ∼17 million reads per library against the human reference genome Hg19, we tracked endogenous DNA content as a function of pre-digestion time. Despite extremely low endogenous DNA content (0.001% to 1.6 %, see Table 1) the pre-digested extracts showed a considerable increase in human DNA (Figure 2), with an average fold-increase of c.2 after 30 minutes and c.3 after 1-6 hours. Despite sample to sample variation, and some extreme outlier values, the average increase of endogenous DNA content observed following pre-digestion was logarithmic (R²=0.87, p=2.5e-5) and seemed to reach the asymptotic maximum after c.1 hour of pre-digestion (Figure 2).

**Figure 2:**
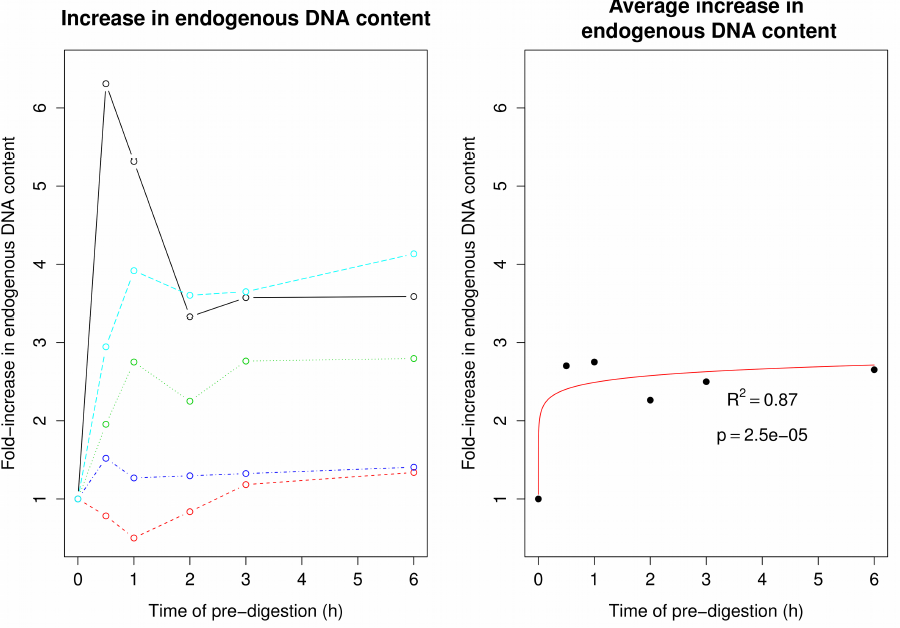
Fold-increase in endogenous DNA content according to pre-digestion lengths. A) Fold-increase in endogenous DNA content in sample EI8 (red), EI9 (green), EI19 (blue), EI22 (cyan), Trinitatis (black). B) A logarithmic model fitted to the mean increase suggests an asymptotic growth (p = 2.5e-5).

Because DNA concentrations decline with longer pre-digestion times, we tested if this decline could be tracked as increased sequence clonality among the human DNA sequences due to endogenous DNA loss. An increase in clonality would result in poor library sequencing efficiency despite a higher endogenous DNA content. Overall, clonality levels were low with average values ranging from 1.49 % (no pre-digestion) to 4.75 % (6 hours of pre-digestion) (see Supplementary Table S1), although we note that library complexity predictions from small datasets of shallow sequencing can give false estimates of high library complexity (Daley and Smith, 2013). Indeed we note that for bone EI8 from which we obtain on average 80 times as many sequences, the clonality increases from 1.3 % (no pre-digestion) to 18 % (6 hours of pre-digestion) causing a complete stagnation in the increase in library efficiency (see Supplementary Table S1).

### Brief pre-digestions on 21 bones

Given that short pre-digestion times appeared to improve access to endogenous DNA content, we next compared the endogenous human DNA content of 21 bones extracted with and without a brief pre-digestion of 15 or 30 minutes (Table 1) in order to confirm the efficiency of the method and determine the statistical significance. The mean enrichment in endogenous DNA content was c. 2-fold (Figure 3), significant as revealed by the one-sided paired t-test results (t = 2.56, df = 20, p-value = 0.009).

**Figure 3:**
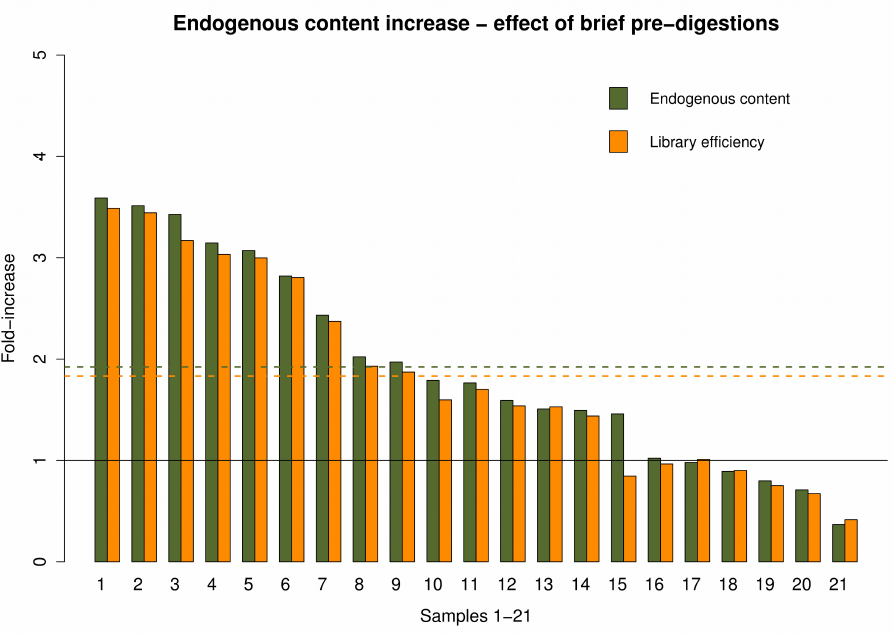
Endogenous content increase following a brief pre-digestion (15-30 minutes). Green colors represent the increase in endogenous content, and yellow colors represents the increase in library efficiency. Dashed lines represent mean values. Values > 1 (black line) are fold-increased and values < 1 are fold-decreased. Samples 1-21 are 1) AAG6, 2) AAG8, 3) ANR 4709, 4) AAG9, 5) AAG7, 6) ANR 4714, 7) AAG2, 8) AAG5, 9) AAG4, 10) Rise 479, 11) ANR 4704, 12) AAG1, 13) AAG3, 14) ANR 4705, 15) Rise 483, 16) ANR 4707, 17) ANR 4708, 18) ANR 4715, 19) ANR 4706, 20) ANR 4710, 21) ANR 4712. The overall increase is significant as revealed by the one-side t-test: (t = 2.56, df = 20, p-value = 0.009).

The overall sequencing efficiency increase was similar to the increase in endogenous DNA content, reflecting that clonality levels were nearly equivalent in the pre-digested and the non-pre-digested sample. Only one sample (Rise483) displayed a clear loss of sequence complexity when pre-digested (Figure 3, Sample 15 and Supplementary Table S2) resulting in lower library efficiency (140353 normalized non-clonal human reads) compared to the non-pre-digested sample (165954 normalized non-clonal human reads).

### Effects of pre-digestion on DNA composition

In general we observed a negligible change in the GC-content in the total datasets (human + non-human) following pre-digestion (Supplementary Fig. S3). The genomic GC-content in the identified human reads was ∼50 % for AmpliTaq Gold amplified libraries and ∼40% GC for Kapa U+ amplified libraries, but there was no correlation between GC-content in the human fraction and pre-digestion times (Supplementary Fig. S2).

Similarly, the DNA damage parameters δs and λ for the human reads displayed no general trend as a function of pre-digestion time and likewise no general pattern was discernable for the decay constants (*k*) and the estimated average fragment length (1/*k*) of the human DNA in the extracts (Supplementary Figure S4-S6, Supplementary Table S4).

### Human DNA from the dentine core and cementum-rich surface of teeth roots

Finally, we investigated whether the cementum-rich outer layer of teeth contained higher endogenous DNA proportions than the dentine, which represents the inner part of the tooth. For 11 of 14 teeth, we observed a higher fraction of human DNA in the cementum compared to the dentine (Figure 4). The mean fold-increase in endogenous DNA proportion was c. 5-fold with values ranging from 0.3-fold to 14-fold. The one-sided paired t-test reveals a significant increase in endogenous DNA in the cementum as compared to the dentine (t = 2.10, df = 13, p-value = 0.05).

**Figure 4:**
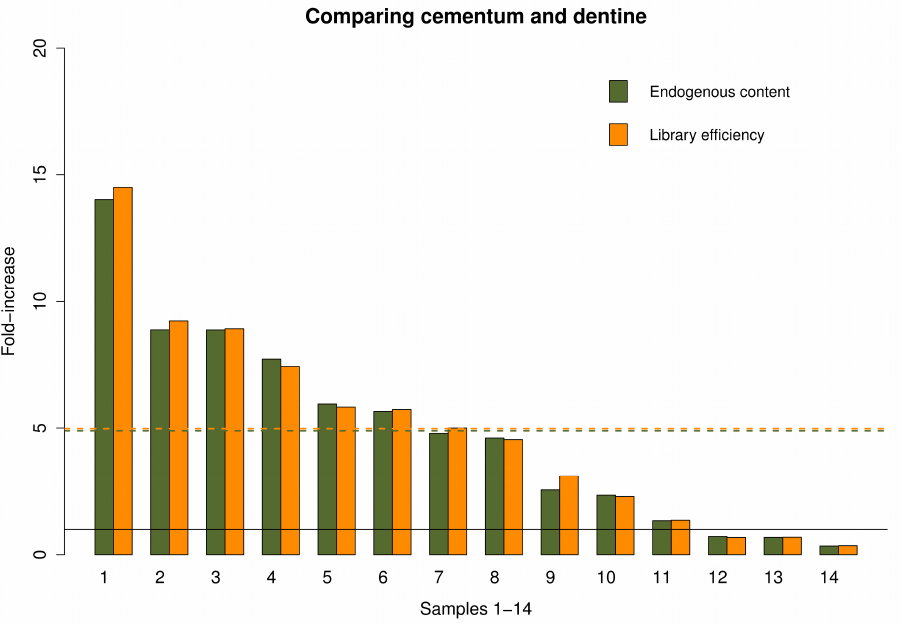
Endogenous content increase in cementum against dentine. Green colors represent endogenous content and yellow colors represent library efficiency. Dashed lines (superimposed) represent mean values. Values > 1 (black line) are fold-increased and values < 1 are fold-decreased. Sample 1-14 are 1) VHM00500 X73, 2) VHM00500 X81, 3) ID-530, 4) VHM00500 X7, 5) VHM00500 X77, 6) ANR 4709, 7) VHM00500 X22, 8) ANR 4711, 9) ID-532, 10) Trinitatis 2, 11) ID-677, 12) ID-678, 13) ANR 4713, 14) Trinitatis 1. The increase is significant as revealed by the one-side t-test: (t = 2.10, df = 13, p-value = 0.05).

**Figure 5:**
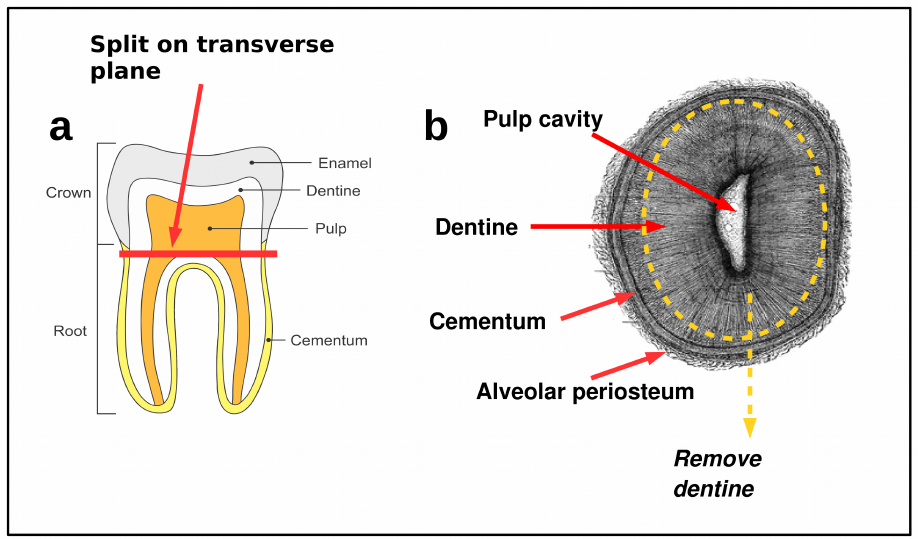
Sampling the cementum enriched hollow roots for aDNA extraction. a) Split the tooth on the transverse plane using round drill-bit, b) Remove dentine from inside the root to create hollow root cap, crush the cementum and digest.

The library efficiency increase was almost identical to the endogenous DNA proportion increase (Figure 4), signifying that extracting from the root surface will generally yield a higher proportion of human DNA without compromising complexity among the template molecules. Finally we note that the cytosine deamination ratio in single-stranded context δs is significantly reduced in 11 out 14 teeth (Table 2).

**Table 2:** Comparing DNA composition parameters for dentine and cementum. Comparison of average length of mapped sequences in dentine and cementum determines that the effect of drilling-induced heating has had little or no impact on the sequence length distribution of the obtained fragments. In contrast, we see significantly increased deamination ratios in dentine compared to cementum as reflected by significantly higher δs means, in 11 out 14 samples. For samples with positive t-value scores on the Welch’s t-test the δs mean in dentine is significantly higher than in the cementum. We cannot determine if the effect is heat-induced or reflects differential preservation.

| | | | Welch's t-test on $\delta s$ distributions from dentine against cementum | | |
| --- | --- | --- | --- | --- | --- |
| Sample number | Substrate | Average length (bp) | T-value | Degrees of freedom | P-value |
| VHM00500 X7_A | Dentine | 61.7 | -10.11 | 56.57 | 2.716E-014 |
| VHM00500 X7_B | Cementum | 60.7 |  |  |  |
| VHM00500 X22_A | Dentine | 64.4 | 11.81 | 56.35 | < 2.2e-16 |
| VHM00500 X22_B | Cementum | 64.6 |  |  |  |
| VHM00500 X73_A | Dentine | 56.4 | 29.79 | 79.29 | < 2.2e-16 |
| VHM00500 X73_B | Cementum | 59.9 |  |  |  |
| VHM00500 X77_A | Dentine | 58.4 | 56.5 | 54.96 | < 2.2e-16 |
| VHM00500 X77_B | Cementum | 64.4 |  |  |  |
| VHM00500 X81_A | Dentine | 72.0 | 397.37 | 48.49 | < 2.2e-16 |
| VHM00500 X81_B | Cementum | 63.4 |  |  |  |
| ANR 4709_A | Dentine | 56.5 | 19.09 | 42.83 | < 2.2e-16 |
| ANR 4709_B | Cementum | 67.9 |  |  |  |
| ANR 4711_A | Dentine | 54.1 | 32.31 | 42.12 | < 2.2e-16 |
| ANR 4711_B | Cementum | 49.7 |  |  |  |
| ANR 4713_A | Dentine | 56.0 | 102.25 | 44.87 | < 2.2e-16 |
| ANR 4713_B | Cementum | 64.6 |  |  |  |
| ID-530_A | Dentine | 56.3 | 153 | 42.82 | < 2.2e-16 |
| ID-530_B | Cementum | 53.9 |  |  |  |
| ID-532_A | Dentine | 48.3 | 3.34 | 40 | 0.001783 |
| ID-532_B | Cementum | 55.3 |  |  |  |
| ID-677_A | Dentine | 61.3 | 12.75 | 43.86 | 2.334E-016 |
| ID-677_B | Cementum | 61.4 |  |  |  |
| ID-678_A | Dentine | 67.3 | 75.03 | 76.27 | < 2.2e-16 |
| ID-678_B | Cementum | 55.7 |  |  |  |
| Trinitatis 1_A | Dentine | 52.6 | -83.28 | 44.47 | < 2.2e-16 |
| Trinitatis 1_B | Cementum | 75.2 |  |  |  |
| Trinitatis 2_A | Dentine | 63.0 | -148.55 | 54.9 | < 2.2e-16 |
| Trinitatis 2_B | Cementum | 63.7 |  |  |  |

### Ancient DNA authentication

For all DNA extractions we observed characteristically elevated δs and λ values (see Supplementary Fig. S6) compared to the reference genome. This suggests that the bulk of the profiled DNA contains templates of ancient origin. The level of DNA contamination was investigated for 12 extracts where we had sufficient data (whole mitochondrial genomes covered with >5X read depth) to meaningfully conduct this analysis (Table 3). Contamination levels proved negligible in 10 extracts confirming that our observed enrichment in endogenous DNA was not driven by modern human contaminant DNA. Encouragingly, in two bone samples, the pre-digestion resulted in a reduced estimated proportion of contamination: from 45% to 1.2% in the Rise479 sample and from 2.5% to 0.3% in Rise483. Likewise, while tooth VHM00500 X81 appeared contaminated in the dentine fraction, contamination levels in the cementum fraction was estimated to be 0.1% (see Table 3 for 95% confidence intervals).

**Table 3:**
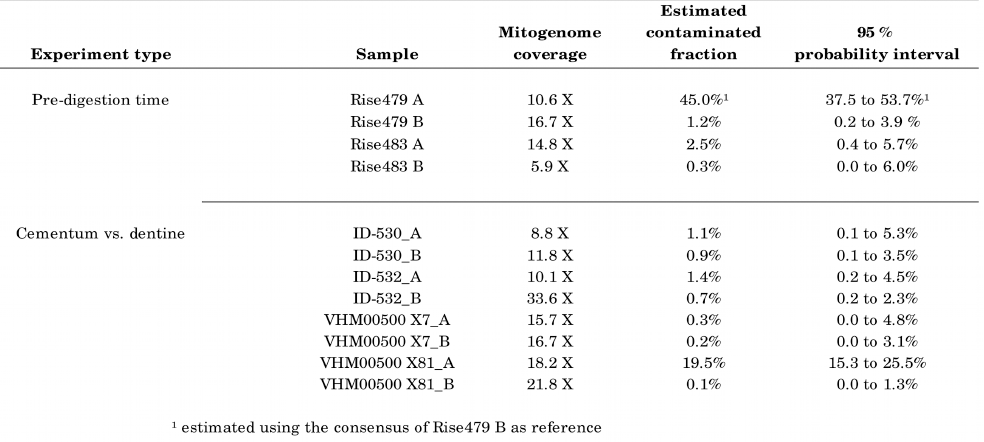
Contamination estimates performed using a Bayesian approach implemented in contamMix. The values represent the estimated fraction of human contaminants. Sample Rise479 A displayed heavy contamination at informative sites (ie. ∼50%) which prevented from determining a consensus sequence with confidence. Instead, Rise479 B (the pre-digested fraction of the bone powder) revealed no signs of contamination. Thus by using the consensus sequence obtained from this extract we were able to estimate the proportion of contaminants in Rise479 A.

| Experiment type | Sample | Mitogenome coverage | Estimated contaminated fraction | 95 % probability interval |
| --- | --- | --- | --- | --- |
| Pre-digestion time | Rise479 A | 10.6 X | 45.0%<sup>1</sup> | 37.5 to 53.7%<sup>1</sup> |
|  | Rise479 B | 16.7 X | 1.2% | 0.2 to 3.9 % |
|  | Rise483 A | 14.8 X | 2.5% | 0.4 to 5.7% |
|  | Rise483 B | 5.9 X | 0.3% | 0.0 to 6.0% |
| Cementum vs. dentine | ID-530_A | 8.8 X | 1.1% | 0.1 to 5.3% |
|  | ID-530_B | 11.8 X | 0.9% | 0.1 to 3.5% |
|  | ID-532_A | 10.1 X | 1.4% | 0.2 to 4.5% |
|  | ID-532_B | 33.6 X | 0.7% | 0.2 to 2.3% |
|  | VHM00500 X7_A | 15.7 X | 0.3% | 0.0 to 4.8% |
|  | VHM00500 X7_B | 16.7 X | 0.2% | 0.0 to 3.1% |
|  | VHM00500 X81_A | 18.2 X | 19.5% | 15.3 to 25.5% |
|  | VHM00500 X81_B | 21.8 X | 0.1% | 0.0 to 1.3% |

<sup>1</sup> estimated using the consensus of Rise479 B as reference

## Discussion

### Pre-digestion time

This study has documented several improvements of immediate value to aDNA research. We show that pre-digestion is a simple and effective means to remove a proportion of non-target DNA from ancient samples. With an average value of 2.7-fold enrichment of the endogenous DNA content following 1 hour of pre-digestion, one would ultimately generate 2.7 times more useable data for the same price.

We interpret the asymptotic increase post-1 hour pre-digestion as a consequence of a gradual change in the ratio of dissolved exogenous DNA over endogenous DNA. This indicates that although the endogenous DNA fraction keeps rising with longer pre-digestions, the benefits of waiting this long are, at best, marginal.

We expected to observe an increase in clonality following longer pre-digestion times because of two complementary phenomena, i) that we have sequenced the human DNA fraction deeper with the same sequencing effort because exogenous DNA is now reduced, and/or ii) that we have sequenced the human fraction deeper because there has been a significant reduction in the DNA library complexity - the human fraction included. The first phenomenon would be a positive outcome of pre-digestion, as it simply requires less sequencing effort before saturation is reached, in a similar manner as described in genomic capture method developments (Carpenter et al., 2013). The second phenomenon would be problematic as it reflects the loss of sequenceable genomic material. As abovementioned, with long pre-digestion times we found near-saturation in the increase of endogenous DNA content, which was accompanied with a loss in DNA concentration (see Supplementary Fig. S1) and, in one case, a considerable increase in sequence clonality. We deduce from these observations that it is advisable to apply this method conservatively using short pre-digestion times (15-30 mins).

We stress that the optimal time of pre-digestion will depend on the temperature used during incubation. Here we have incubated at 50°C, but if incubating at lower temperatures, it may be advantageous to apply a longer pre-digestion step.

### Changes in genomic composition and DNA damage

It is conceivable that prolonged incubation times at 50°C could result in increased DNA damage compared to non-pre-digested samples. However, for the pre-digestion lengths tested in this study, we found no general trends on the estimated DNA damage parameters (see Supplementary Fig. S4-S6 and Supplementary Table S4 for damage parameters, Supplementary Fig. S2 for GC-content). This observation provides evidence that the pre-digestion procedure will not damage the DNA or otherwise alter the genomic composition in the extract compared to non-pre-digested samples. This could be because the pre-digestions used in these experiments are relatively brief (< 6h hours) and unspecific (ie. dissolving both the organic and mineral phases of the samples) and hence they do not provide the basis for comparing human DNA preserved within different subniches, such as that anlyzed in (Schwarz et al., 2009).

### Comparing the dentine core and cementum-rich surface of tooth roots

Using n=14, we observe a significant increase (one-sided t-test: p = 0.05), in endogenous DNA content when extracting DNA from the hollow root cap compared to the drilled-out dentine. This demonstrates that cementum is a highly advantageous substrate for aDNA extractions. Previous studies have estimated that the cementum-rich surface contains five times higher amounts of human mtDNA than dentine (Adler et al., 2011). However, these higher concentrations of human DNA are only relevant for shotgun sequencing if they are also manifested in a higher *proportion* of human DNA. Our results demonstrate that this is indeed the case, and this is crucial knowledge when preparing a tooth for shotgun DNA sequencing. We note that three outliers in which the endogenous DNA fraction is decreased likely represents deviations from the general preservation conditions due to extrinsic factors, such as broken or punctured roots which we intuitively deem as advisable to consider as a criteria when sampling teeth for DNA extractions.

The contamination analyses (Table 3) rejected that the elevated fraction of human DNA in the outer root layers was simply due to modern human contamination. Hence, when teeth are available for aDNA studies, the results presented herein strongly support targeting the cementum-rich root surface, if possible. As a side note, using only the root of a tooth for the aDNA extraction facilitates leaving the crown intact, which can then be used for morphological investigations.

## Recommendations and conclusion

We recommend following these five points when extracting aDNA from ancient bone or teeth:

1. Apply a brief pre-digestion step (15-30 mins).
2. If sufficient material is available then run several extractions in parallel, possibly with differing pre-digestion lengths, adequately adjusted to the sample amount and preservation state.
3. Do not discard the supernatant (the pre-digest) at first, as it will contain a fraction of endogenous DNA molecules and could become useful, in particular if the sample contains a high endogenous DNA proportion to start with.
4. Do not discard undigested bone pellets post-24-hour digestion, as they should contain a higher endogenous DNA fraction than the first extraction (whether this was pre-digested or not).
5. Always sample the cementum-rich surface of tooth roots in favor of dentine but be careful of potential surface contamination (ie. scrape off the outermost periosteum layer with a scalpel or a drill).

We advocate caution when implementing pre-digestions, especially if only very little starting material is available (i.e. <50mg). In such case it may not be recommendable to pre-digest the sample because the DNA concentration in the final extract may become critically low. Finally, we note that if pre-digestion is combined with other methods that have demonstrated an enrichment in the endogenous DNA fraction, such as the capture of mitogenomes (Maricic et al., 2010) or whole genomes (Carpenter et al., 2013), single-stranded sequencing libraries (Gansauge and Meyer, 2013; Meyer et al., 2013), or damage enriched single-stranded sequencing libraries (Gansauge and Meyer, 2014), it can likely result in a many-fold increase in endogenous DNA proportions. We believe that implementing these inexpensive procedures will increase the discovery rate of samples suitable for ancient genomic research considerably and could prove useful for any discipline working with degraded DNA from biological remains, such as forensic sciences.

## Acknowledgments

We thank our colleagues Anna-Sapfo Malaspinas, José Victor Moreno-Mayar, Morten Rasmussen, Thorfinn Korneliussen, Lasse Vinner, and Peter Ilsøe for insightful discussions, and the Danish National High-throughput DNA Sequencing Centre and Jesper Stenderup for technical assistance. We are grateful to Erik Thorsby (Dept. of Immunology, University of Oslo), Reidar Solsvik (The Kon-Tiki Museum), and Per Holck (The Anatomical Institute, University of Oslo) for access to the Easter Island samples. We thank Sidsel Wåhlin (The Historical Museum of Vendsyssel) for the Danish Iron Age samples, Jette Arneborg (The Greenland National Museum and Archives) and Niels Lynnnerup (Dept. of Forensic Medicine, University of Copenhagen) for the Greenlandic samples, Jacob Mosekilde (Museum of Copenhagen) for the Danish Trinitatis samples, and Menno Hoogland (Leiden University) for the samples from Guadeloupe. The Bronze Age samples from Hungary were collected and analysed as part of “The Rise” project funded by the European Research Council (FP/2007-2013, grant no. 269442 to Kristian Kristiansen). The Danish and Greenlandic samples (Trinitatis, Iron Age, and Norse Eastern Settlement in Greenland) were collected and analysed as part of “The Genomic History of Denmark” project (KU2016) funded by the University of Copenhagen. HS was supported by a Synergy Grant from the European Research Council (FP7/2007-2013, grant agreement no. 319209). PBD was supported in part by the Siemens Foundation, and MEA was supported by the Marie Curie Actions of the European Union (FP7/2007-2013, grant no. 300554). GeoGenetics is supported by the Lundbeck Foundation and the Danish National Research Foundation.

## Author Contributions

MEA and EW designed the study. PBD carried out the lab work with assistance from MEA, HS and AM. PBD and MEA analyzed the data and wrote the manuscript, with considerable input from LO and the remaining authors.

